# Defining of MAbs-neutralizing sites on the surface glycoproteins Gn and Gc of a hantavirus using vesicular stomatitis virus pseudotypes and site-directed mutagenesis

**DOI:** 10.1101/307975

**Authors:** Lev Levanov, Rommel Paneth Iheozor-Ejiofor, Åke Lundkvist, Olli Vapalahti, Alexander Plyusnin

**Author notes:** Address correspondence to Lev Levanov. These authors contributed equally to this study (Lev Levanov, Rommel Paneth Iheozor-Ejiofor). Åke Lundkvist. Olli Vapalahti –.

## Abstract

Earlier four Monoclonal antibodies (MAbs) against surface glycoproteins Gn and Gc of Puumala virus (PUUV, genus *Orthohantavirus*, family *Hantaviridae*, order *Bunyavirales*) were generated and for three MAbs with neutralizing capacity the localization of binding epitopes was predicted using pepscan and phage-display techniques. In this work, we produced vesicular stomatitis virus (VSV) particles pseudotyped with the Gn and Gc glycoproteins of PUUV and applied site-directed mutagenesis to dissect the structure of neutralizing epitopes. Replacement of cysteine amino acid (aa) residues with alanines resulted in pseudotype particles with diminished (16 to 18%) neut-titers; double Cys→Ala mutants, as well as mutants with bulky aromatic and charged residues replaced with alanines, have shown even stronger reduction in neut-titers (from 25% to the escape phenotype). *In silico* modelling of the neut-epitopes supported the hypothesis that these critical residues are located on the surface of viral glycoprotein molecules and thus can be recognized by the antibodies indeed. Similar pattern was observed in experiments with mutant pseudotypes and sera collected from patients suggesting that these neut-epitopes are utilized in a course of human PUUV infection.

**IMPORTANCE:** Neutralization of viruses by antibodies is one of the key events in infection. We identified a set of mutations in the surface proteins of PUUV that reduced the virus-neutralizing activity of MAbs. Moreover, we found three mutants with escaped phenotype. These data will help understanding the mechanisms of hantavirus neutralization and assist construction of vaccine candidates.

## INTRODUCTION

Puumala virus (PUUV) is a typical zoonotic agent. It causes the mild form of haemorrhagic fewer with renal syndrome, also called Nephropathia epidemica (NE), occurring in most of Europe where its rodent host, the bank vole (*Myodes glareolus*) prevails. Currently there are few inactivated hantavirus vaccines licensed in East-Asian countries (1, 2) and no hantavirus vaccine approved for use outside this region. A phase 2 study on a human PUUV/Haantan virus (HTNV) DNA vaccine is ongoing (1, 2, 3).

PUUV belongs to *Orthohantavirus* genus in the family *Hantaviridae*, order *Bunyavirales*. The negative-sense RNA genome of hantaviruses is divided into small (S), medium (M) and large (L) segments, which encode respectively for the nucleocapsid (N) protein (in some hantaviruses, also NSs protein), glycoprotein precursor processed into the surface Gn and Gc glycoproteins, and RNA-dependent RNA polymerase (L-protein). The genome RNA segments are encapsidated by the N protein within the virion. The virions are commonly spherical and vary in size from 120 to 160 nm (4, 5, 6, 7). They comprise a lipid envelope covered with tetrameric Gn-Gc spike compexes (8, 9, 10). The glycoproteins carry major immunodominant and structural epitopes. It is thought that they bind receptor(s) on the surface of the target cells and mediate the fusion of viral and cellular membranes (11). They also induce virus-neutralizing antibodies responsible for protective immune response (12, 13, 14).

Neutralization of viruses by antibodies is one of the key events in infection. Good understanding of the mechanisms of neutralization and the structure of epitopes are essential for studying immune response and development of effective diagnostics and vaccines. Earlier the epitopes for two bank vole and one human neutralizing monoclonal antibodies (MAbs) against PUUV were identified using pepscan and phage display techniques (15). In this study, we created vesicular stomatitis virus (VSV) particles pseudotyped with the surface Gn and Gc glycoproteins of PUUV and applied site-directed mutagenesis to dissect the structure of neutralizing epitopes.

## RESULTS

### *In silico* modelling of initial and mutant glycoproteins of PUUV

Earlier the binding sites for MAbs 5A2, 1C9 and 4G2 against PUUV Gn and Gc glycoproteins were predicted using pepscan and phage-display. The most reactive sites were considered potential neutralizing epitopes (15). Two binding sites were found for MAb 5A2 (epitope A: 61SLKLESSCNFDL72 and epitope B: 264EPLYVPTLDDYRSAEVL280), one binding site was found for Mab 1C9: 822EQTCKTVDSNDCL834 and one site for MAb 4G2: 904KCAFATTPVCQFDGNTIS921. The sites for MAb 5A2 are located in the Gn glycoprotein and the sites for the other two MAbs in the Gc glycoprotein (Fig. 1). Sequences of the epitopes are highly conserved within known PUUV genetic lineages; moreover many aa residues are conserved also in closely related hantavirus species like Tula and Sin Nombre viruses (Fig. 1).

**Figure 1.**
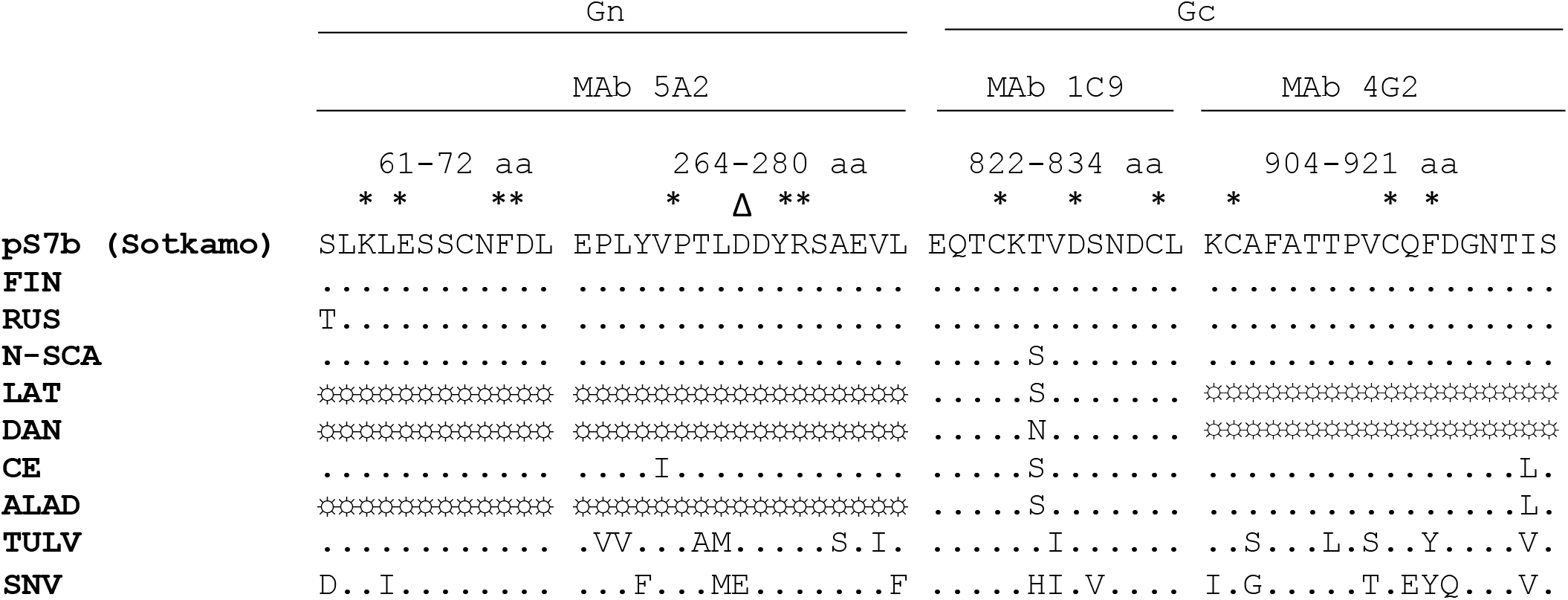
Epitopes on Gn and Gc glycoproteins of PUUV recognized by MAbs: aa sequence alignment of different genetic lineages. FIN, Finnish lineage (strain Pieksämäki); RUS, Russian lineage (strain Kazan); N-SCA, North Scandinavian lineage (strain Umeå); LAT, Latvian lineage (strain Latvia 149); DAN, Danish lineage (strain Fyn47); CE, Central European lineage (strain Ardennes 75); ALAD, Alpe-Adrian lineage (strain Croatia/Gerovo 982). In addition, corresponding regions of Tula virus (TULV, strain TulaV02) and Sin Nombre virus (SNV, strain NM H10) are shown. Aa residues replaced with alanines are marked with asterisks; the aa residue replaced with valine is marked with a triangle; not available sequences are marked with suns.

In the mutagenesis experiments, we targeted conserved cysteines, bulky aromatic (Y, F) and charged (K, R, D, E) aa residues; one proline residue was also mutated. These aa residues could be critical for maintaining the proper fold of a polypeptide chain, likely in close interaction with a membrane. Charges also can play a role in long-distance interactions, e.g. the interactions between epitopes and neutralizing antibodies.

To visualize the epitopes on Gn and Gc glycoproteins and evaluate possible effect of mutations on their structure, Yasara software and pdb-structures based on crystal structures of PUUV Gn and Gc glycoproteins in dimeric and monomeric form respectively were used (Table 1). It appeared that all epitopes for MAbs were located on the surface of Gn and Gc glycoproteins and hence available for corresponding MAbs (Table 1). Replacement of conservative aromatic aa residues with alanines in the epitopes for MAbs 5A2 (F70A, Y274A) and 4G2 (F915A) resulted in visible structural changes (Table 1). Similar results were obtained when lysine for epitope A (K63A) and proline for epitope B (P269A) of Mab 5A2 were replaced with alanines. Structural changes of the epitopes induced by other mutations were less pronounced (Table 1).

**Table 1.**
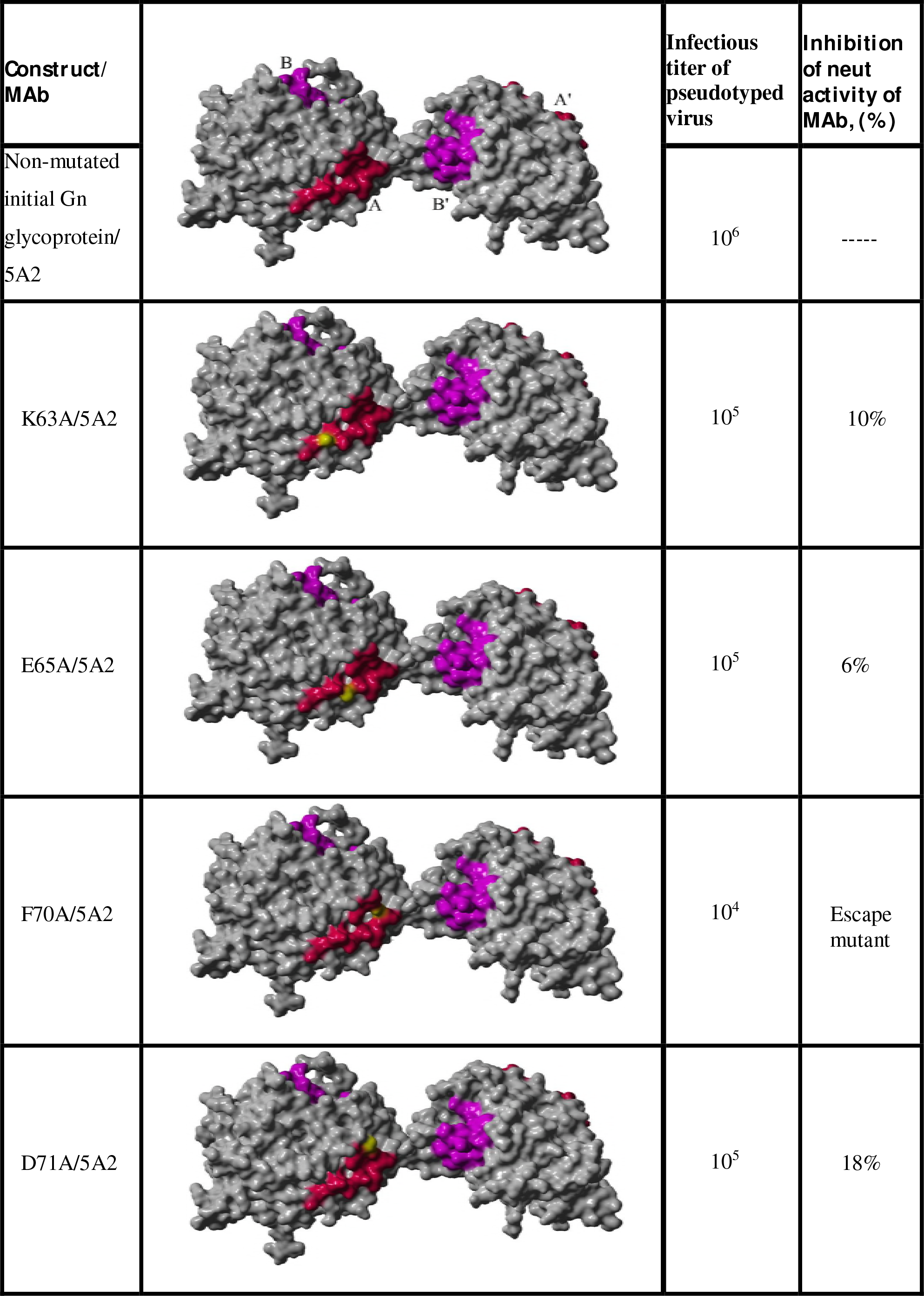
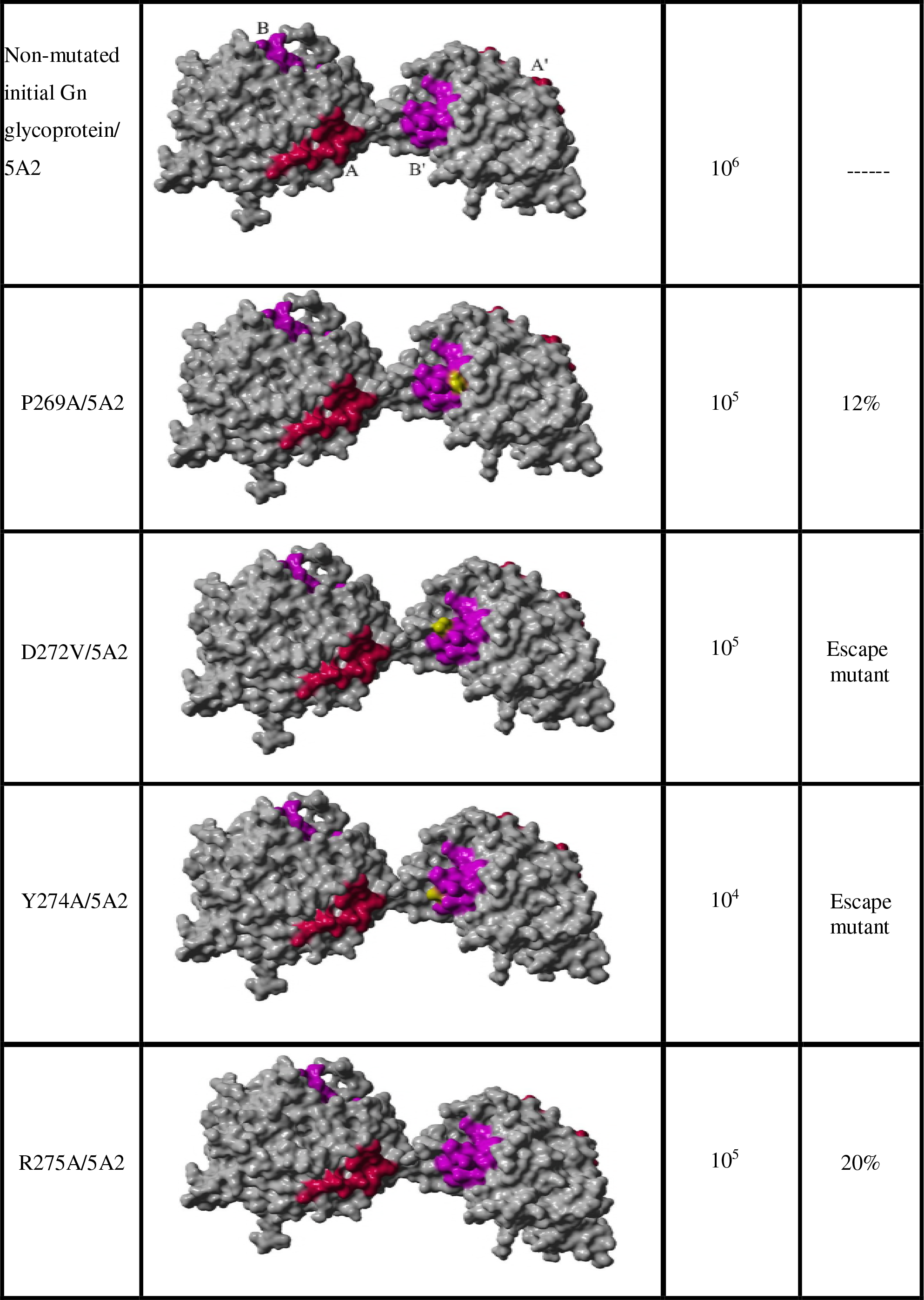
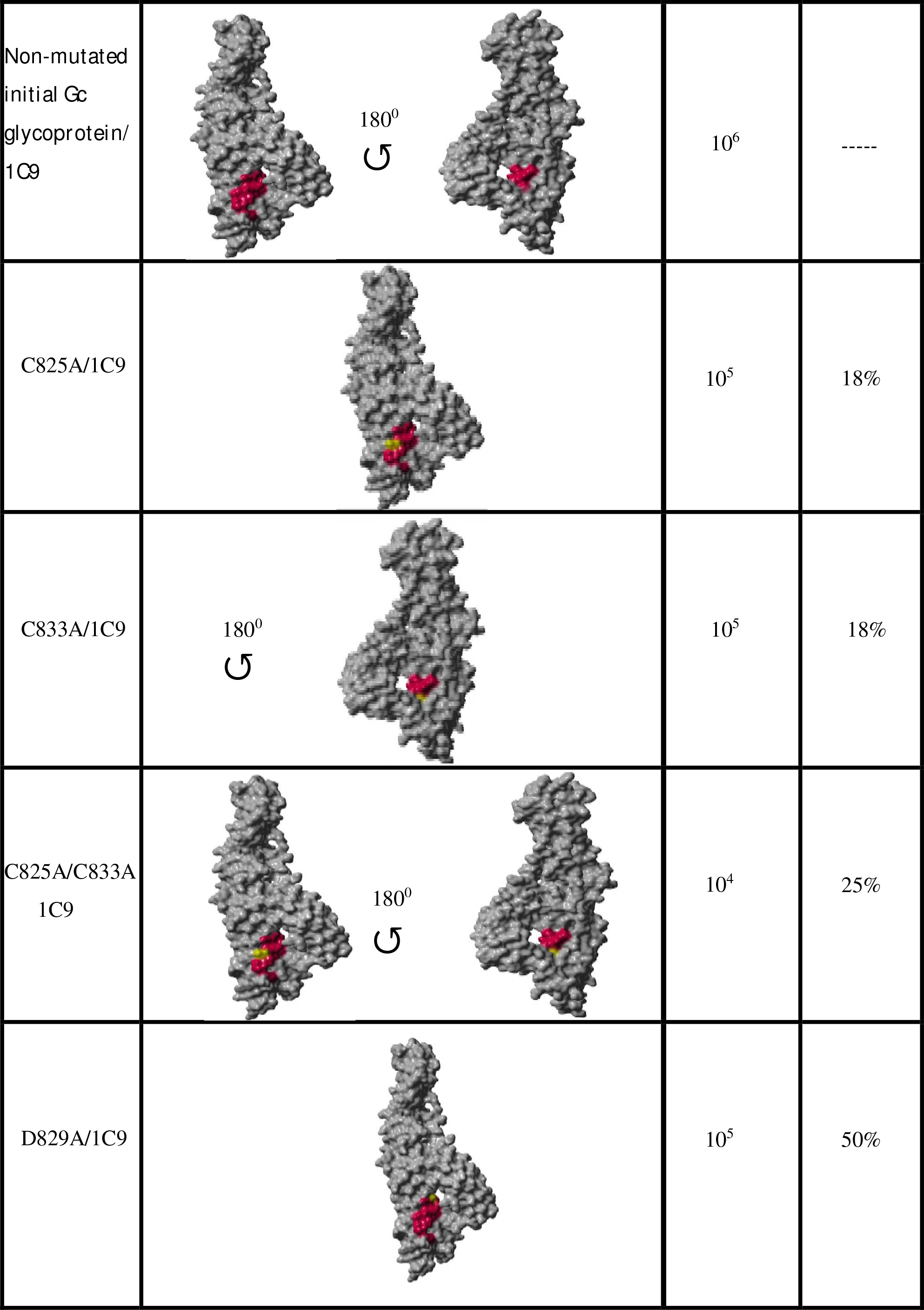
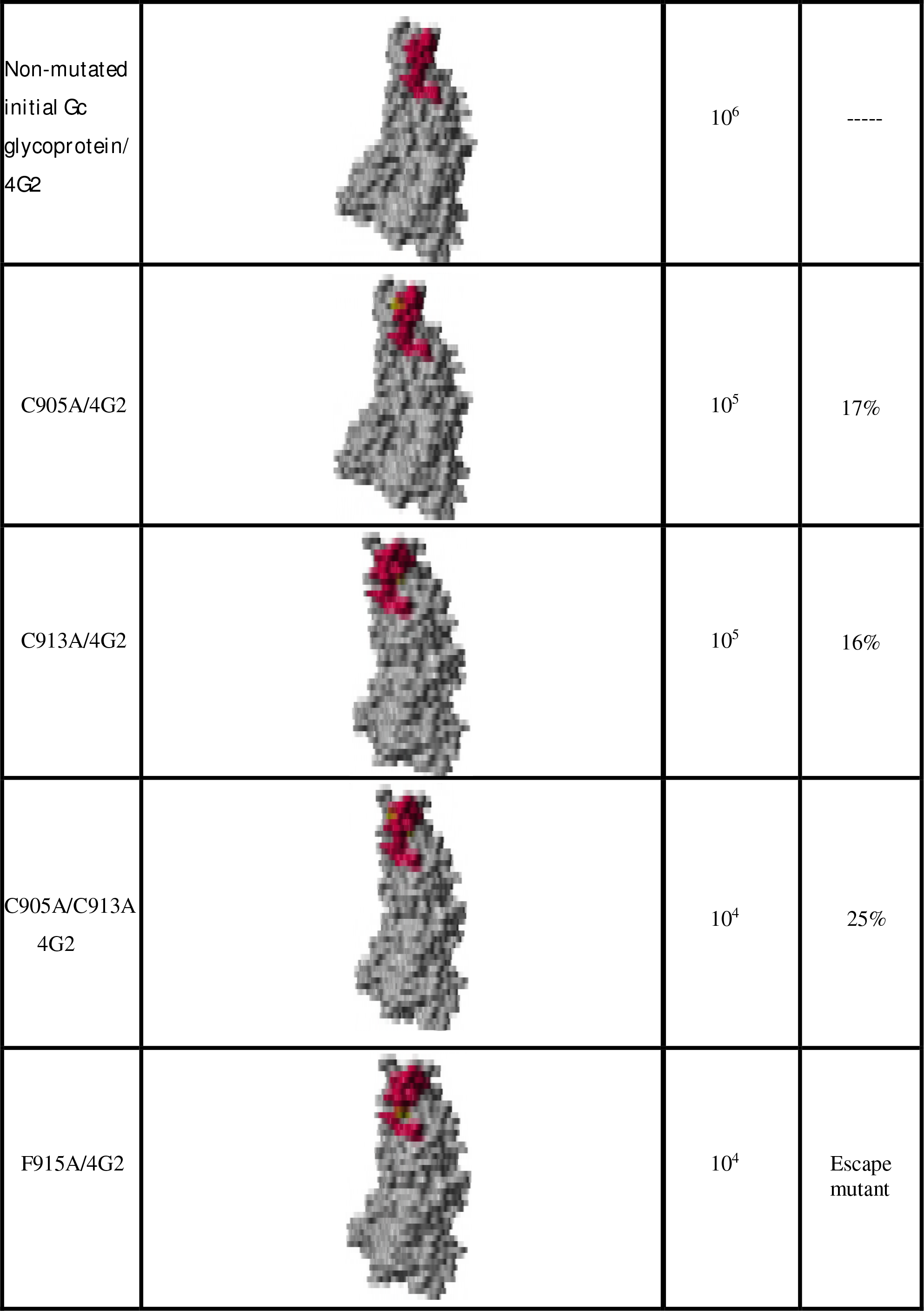
Models of Gn and Gc glycoproteins of PUUV (YASARA software), infectious titers of VSV particles pseudotyped with the corresponding Gn and Gc glycoproteins and MAb activities in neutralization assay with respective pseudotypes. Gn glycoprotein is represented as a dimer, Gc glycoprotein is represented as a monomer. The color code: magenta - neutralizing epitopes for MAbs 1C9, 4G2 and epitopes A, Aʹ-for Mab 5A2; purple - epitopes B and Bʹ for Mab 5A2; yellow – mutated aa residues.

| Construct/<br>MAb |  | Infectious<br>titer of<br>pseudotyped<br>virus | Inhibition<br>of neut<br>activity of<br>MAb, (%) |
| --- | --- | --- | --- |
| Non-mutated<br>initial Gn<br>glycoprotein/<br>5A2 | | $10^6$ | ----- |
| K63A/5A2 | | $10^5$ | 10% |
| E65A/5A2 | | $10^5$ | 6% |
| F70A/5A2 | | $10^4$ | Escape<br>mutant |
| D71A/5A2 | | $10^5$ | 18% |
| Non-mutated<br>initial Gn<br>glycoprotein/<br>5A2 | | $10^6$ | ----- |
| P269A/5A2 | | $10^5$ | 12% |
| D272V/5A2 | | $10^5$ | Escape<br>mutant |
| Y274A/5A2 | | $10^4$ | Escape<br>mutant |
| R275A/5A2 | | $10^5$ | 20% |
| Non-mutated<br>initial Gc<br>glycoprotein/<br>1C9 | | $10^6$ | ----- |
| C825A/1C9 | | $10^5$ | 18% |
| C833A/1C9 | | $10^5$ | 18% |
| C825A/C833A<br>1C9 | | $10^4$ | 25% |
| D829A/1C9 | | $10^5$ | 50% |
| Non-mutated<br>initial Gc<br>glycoprotein/<br>4G2 | | $10^6$ | ----- |
| C905A/4G2 | | $10^5$ | 17% |
| C913A/4G2 | | $10^5$ | 16% |
| C905A/C913A<br>4G2 | | $10^4$ | 25% |
| F915A/4G2 | | $10^4$ | Escape<br>mutant |

### Construction and expression of mutant Gn and Gc glycoproteins of PUUV

To study the impact of aa substitutions described above on the expression and folding of PUUV glycoproteins, the respective constructs were produced and transfected into 293T cells. When the cells were lysed and immunoblotted with rabbit polyclonal anti-PUUV Gn and anti-PUUV Gc sera, both Gn and Cc glycoproteins were detected (data not shown). The characterized constructs were then used for pseudotyping of VSV particles.

### Pseudotyping of rVSVΔG EGFP with mutant Gn-Gc glycoproteins of PUUV

To pseudotype VSV particles with mutant Gn and Gc glycoproteins, the corresponding mutant plasmids were transfected into 293T cells. Following 48h, the transfected cells were infected with rVSVΔG EGFP. The supernatant was harvested after 48h, ultracentrifugated and titrated on Vero E6 cells. It was shown that all 16 mutant glycoproteins were incorporated into rVSV particles. The titers for the pseudovirion mutants with aromatic and double cysteine mutants were two logs lower (10^4^), compare to the pseudotypes obtained from initial control plasmid pS7b (10^6^). Mutants with only one cysteine substitution, proline substitution and those in which charged amino acids were replaced with alanines could be titrated up to 10^−5^.

### Neutralization studies

The effect of different aa substitutions on the binding and neutralizing properties of mutant pseudotyped VSV particles was analyzed with MAbs (Fig. 2) and human serum samples positive for PUUV in neutralization assay.

**Figure 2.**
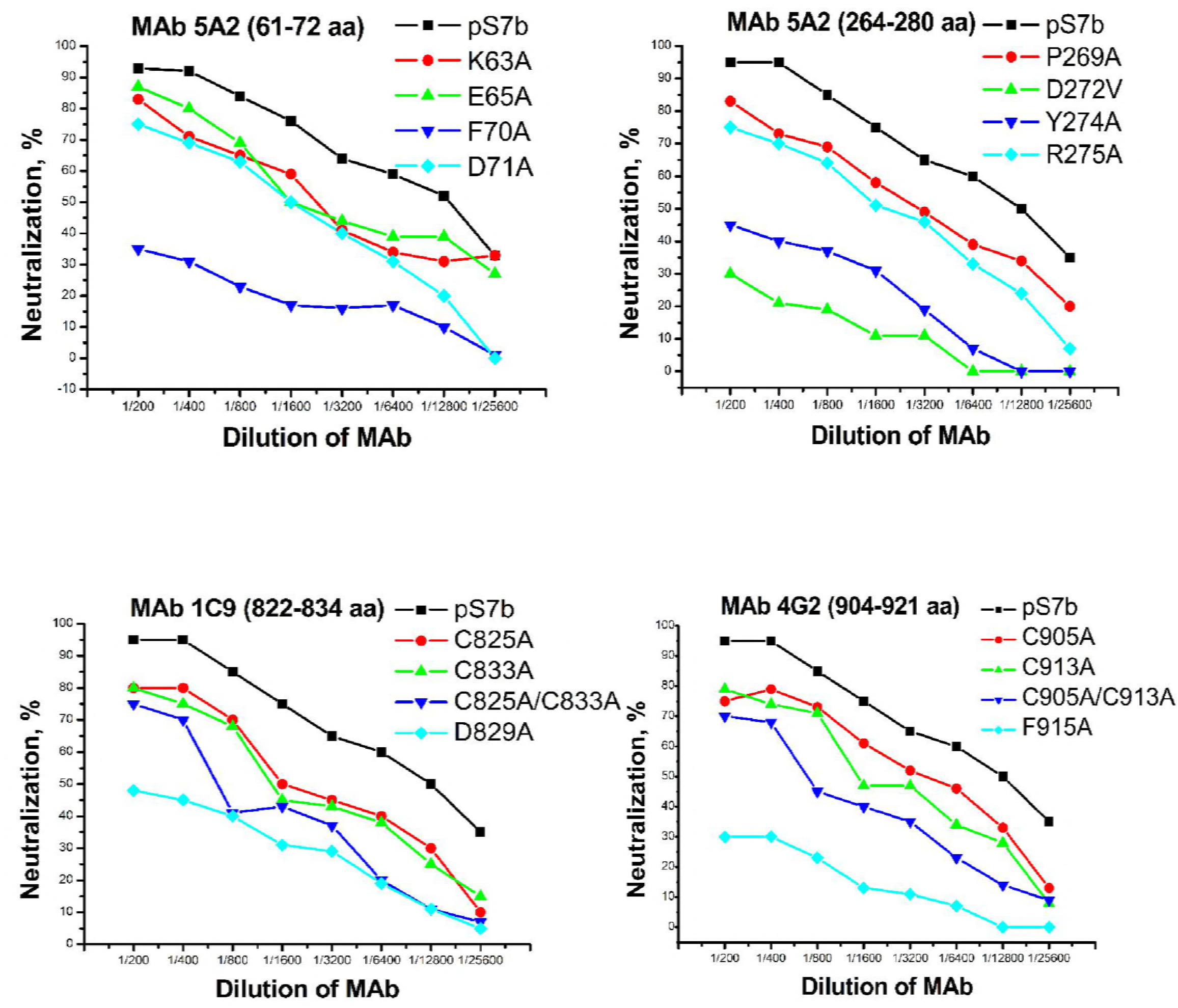
Neutralizing activity of MAbs against PUUV with VSV particles pseudotyped with mutant PUUV Gn and Gc glycoproteins.

It was shown that replacement of conservative aromatic aa residues with alanines in the epitopes for MAbs 5A2 (F70A, Y274A) and 4G2 (F915A) resulted in escape phenotypes of the corresponding pseudotype viruses. Similar results were obtained for the mutant pseudotype virus D272V in neutralization assay with Mab 5A2 (Table 1).

The replacement of charged amino acids in epitope A: K63A, E65A, D71A resulted in a loss of neutralizing activity of MAb 5A2 by 10%, 6% and 18%, respectively (Table 1). Pseudotyped viruses with mutations P269A and R275A showed diminished binding properties of MAb 5A2 by 12% and 20%, respectively (Table 1).

The neutralizing activity of MAb 1C9 was dropped by 18% in the presence of pseudotyped viruses C825A and C833A and by 25% in the presence of double C825A/C833A mutant. Substitution D829A provided 50% protection of the pseudotyped virus from neutralization by MAb 1C9 (Table 1).

The C905A, C913A and C905A/C913A mutations diminished neutralizing properties of Mab 4G2 by 17%, 16% and 25%, respectively (Table 1).

All mutants described above showed somewhat diminished neut-titers of positive human NE-serum samples; the reduction however was not statistically significant.

## DISCUSSION

In this study, we created VSV particles pseudotyped with the surface Gn and Gc glycoproteins of PUUV and applied site-directed mutagenesis to dissect the structure of neutralizing epitopes for MAbs 5A2, 4G2 and 1C9. It is generally agreed that epitopes are enriched in charged and polar aa residues and depleted of aliphatic hydrophobic residues (16, 17, 18). They may contain cysteines which can expose other amino acids on the surface and form epitope utilizing disulfide bonds (16). It was suggested that the tyrosine and tryptophan residues are significantly over-presented on the surface of epitopes and that valine is significantly depleted (19). In our study, *in silico* modelling demonstrated pronounced structural changes in epitopes for those mutant proteins in which aromatic amino acids were replaced with alanines. These substitutions also dramatically diminished neutralizing capacity of MAbs in neutralization assay with mutant pseudotypes. Structural changes in epitopes with replaced charged amino acids were less strong. These mutant VSV pseudotypes showed diminished biological properties of MAbs as well.

Interestingly, the mutation D272V described earlier for PUUV escape mutant (20) had the same effect on the phenotype of pseudotyped virus in neutralization assay with Mab 5A2. According to multimeric model for PUUV Gn glycoprotein on the surface of virion offered by Li et al (21), the D272 is located roughly in the middle of contact area between two epitopes (A and B^/^) of two neighboring monomeric subunits of Gn glycoprotein.

The mutant constructs in which only one cysteine in Gc glycoprotein was replaced with alanine demonstrated moderate reduction of titers in neutralization assay compare to aromatic and charged amino acid replacements. The most significant reduction of neut-titers was found when both cysteines were substituted with alanines; the cumulative effect of these two mutations was observed.

Importantly, reduced neutralizing titers were observed when all mutant constructs were tested with human serum samples positive for PUUV, even if this reduction was not statistically significant. One explanation could be that positive sera recognize different epitopes on the surface of Gn and Gc glycoproteins of PUUV. Earlier, it was demonstrated by pepscan, that human serum samples positive for PUUV antibodies were bound to aa residues 19-33, 52-72, 79-93 and 85-99 of Gn glycoprotein and aa residues 442-453, 946-957 and 955-966 of Gc glycoprotein (15).

In conclusion, we analyzed putative MAb-neutralizing epitopes on PUUV Gn and Gc glycoproteins. Since “classical”reverse genetics, including the virus rescue from plasmids, is not available for hantaviruses yet, the glycoproteins were mutated using VSV-pseudotypes. Altogether 14 point and two double mutations were introduced into Gn and Gc molecules and all 16 recombinant viruses were successfully rescued. Analysis of infectious and neutralizing titers of mutant viruses confirmed that the MAb-neutralizing epitopes are located in the regions suggested in earlier studies which used pepscan and phage-display techniques. Mutagenesis revealed an importance of conserved bulky aromatics (F, Y) and charged (K/R, E/D) aa residues, as well as cysteines and perhaps also prolines, for maintaining the proper structure of the epitopes. Most notably, in addition to D272V mutation that was earlier found responsible for changing of PUUV phenotype to escape-type (20), we were able to generate three more escape mutants, all three by replacing a single bulky-aromatic aa residues: F70A, Y274A and F915A. 3D-modelling of the presumable MAb neutralizing epitopes was proven very helpful and confirmed e.g. that the epitopes are located on the surface of the Gn and Gc molecules and hence are available for antibody recognition.

The newly gained knowledge will aid in preparing more efficient diagnostics and vaccines.

## MATERIALS AND METHODS

### Construction of the plasmids encoding mutant Gn-Gc glycoproteins of PUUV

To introduce mutations into Gn and Gc glycoproteins of PUUV, modified pS7b construct (22) was used. This fully synthetic and codon-optimized construct is based on the M segment of PUUV, strain Sotkamo (23) and has one additional amino acid (aa) substitution (G822E) compare to the parental strain. The substitution was likely appeared during early passages of the original PUUV isolate in Vero E6 cells. Point-directed mutagenesis was accomplished using Phusion site-directed mutagenesis kit (Thermo Fisher Scientific). Altogether 14 single and two double mutations were introduced and the corresponding pseudotyped VSV particles rescued.

### Rescue of recombinant VSVΔG EGFP (rVSVΔG EGFP)

The detailed rescue protocol of rVSVΔG EGFP was described earlier (22). The basic plasmid carries the genome of VSV, strain Indiana, in which the G protein-encoding gene is replaced with the enhanced green fluorescent protein (EGFP). The basic plasmid encoding the VSVΔG EGFP antigenomic RNA, together with the helper plasmids expressing the viral N, P, L and G proteins were co-transfected into BSRT7 cells. After 48h incubation, the supernatant was collected, clarified and used for amplification of recombinant VSV stock on the next step. To amplify the viral stock, 293T cells transfected with the plasmid encoding glycoprotein G of VSV, were infected with the rVSVΔG EGFP from the previous step. After 48h, supernatant was collected, clarified, pelleted through a 30% sucrose cushion, resuspended in PBS and frozen at - 80°C until further use.

### Pseudotyping of rVSVΔG EGFP with mutant Gn-Gc glycoproteins of PUUV

At 48h after transfection of 293T cells with the mutant constructs or control plasmids (pS7b and pCAGGS/MCS), the cells were infected with rVSVΔG EGFP at a multiplicity of infection of 1.0 for 1h at 37°C. The transfected-infected cells were then washed three times with 1% heat-inactivated FCS-PBS, and culture medium was added. After 48h, the culture supernatants were clarified by centrifugation, pelleted through a 30% sucrose cushion and resuspended in PBS. All pseudovirion stocks were frozen at −80°C until use. Titration of the rVSVΔG EGFP and pseudotype virions was done as described previously (24), and the foci were counted using a fluorescent microscope.

### Neutralization assay

A total of 50 μl of medium containing 150 fluorescence focus forming units of VSV pseudotypes was incubated with an equal volume of serially diluted MAb or serum sample for 1h at 37°C. Then 90 μl of the mixture was inoculated onto Vero E6 cells monolayers in 96-well tissue culture plates. After adsorption for 1h, the inoculum was replaced with Eagle minimum essential medium. After 18 h, cells infected with fluorescent VSV pseudotypes were examined and counted.

### 3D-modelling

Three-dimensional models for initial and mutated PUUV Gn and Gc glycoproteins were generated based on known crystal structures (25, 26); pdb code 5fyn for Gn glycoproteins in dimeric form and pdb code 5j81.1.A for Gc glycoprotein in monomeric form, using Yasara software. The models were ranked on the basis of alignment score (PSI_BLAST) and structural quality (Z_Score). The models generated had following QMEAN6 and Z_Score intervals: 0.711-0.715 and −1.500–1.564 for epitope the A of MAb 5A2; 0.710-0.718 and – 1.422-1.614 for the epitope B of MAb 5A2; 0.680-0.683 and – 2.233-2.325 for the epitope of MAb 1C9; 0.676-0.684 and – 2.212-2.417 for the epitope of MAb 4G2. Based on these parameters, the models were considered accurate for visualization of epitopes.

## ACKNOWLEDGMENTS

This study was funded by the Academy of Finland, Finnish Cultural Foundation and Juhani Ahon Medical Research Foundation sr. We thank Dr. John Rose (Yale University) for VSV rescue plasmids. Our gratitude also goes to Dr. Jussi Hepojoki (University of Helsinki) for helpful discussion. We appreciate an excellent technical assistance of Irina Suomalainen.

